# Apical clathrin-coated pits control the location, timing, and scale of microvillar growth

**DOI:** 10.1101/2024.10.21.619431

**Authors:** Olivia L. Perkins, Alexandra G. Mulligan, Evan S. Krystofiak, K. Elkie Peebles, Rekha N. Nagarajan, Leslie M. Meenderink, Bryan A. Millis, Matthew J. Tyska

## Abstract

Epithelial cells from diverse contexts assemble apical specializations to serve tissue-specific functions. In virtually all cases, these features consist of arrays of microvilli: micron-scale, actin bundle-supported protrusions that mediate biochemical and physical interactions with the external environment. Despite their importance for epithelial physiology, how microvilli grow during cellular differentiation remains poorly understood. Using genetic and small molecule perturbations, we found that an epithelial cell’s potential for growing microvilli of normal size is limited by an adjacent actin-dependent process: apical clathrin-mediated endocytosis. Unexpectedly, timelapse imaging of individual microvillar growth events revealed tight spatial and temporal coupling to sites of clathrin-mediated endocytosis. Ultrastructural characterization of undifferentiated epithelial monolayers also showed that most nascent microvilli are in contact with an apical endocytic pit. Finally, inhibition of the Arp2/3 branched nucleation complex, which drives actin polymerization on coated pits, significantly reduced the accumulation of new microvilli on the surface of differentiating epithelial cells. Based on these discoveries, we conclude that clathrin-mediated endocytosis and its associated Arp2/3-based actin nucleation activity control the timing and location of microvillar growth, as well as the dimensions of the resulting protrusions.

## INTRODUCTION

Actin-based membrane protrusions extend from the surface of virtually all animal cells, where they mediate biochemical and physical interactions with the extracellular environment. Microvilli are canonical examples; as finger-like protrusions that are found in large arrays on the apical surfaces of solute transporting epithelial tissues, they significantly increase the plasma membrane surface area available for holding the channels and transporters that are central to physiological function ^1, 2^. Variations on this morphological theme are found throughout the animal kingdom, on organisms ranging from ancient single-celled protists to mammals ^3, 4^. Interestingly, previous evolutionary studies suggested that as mammalian cell functions became more specialized, building arrays of microvilli enabled epithelial cells to increase surface area for maximizing solute uptake potential ^3, 5^. In specific tissues, such as the intestinal tract and kidney tubules, epithelial cells build highly ordered arrays of microvilli known as brush borders, where the apical surface is nearly saturated with protrusions, characterized by hexagonal spacing when viewed *en face* ^6^.

The ultrastructure and proteome of intestinal microvilli have been well characterized in studies spanning decades ^7, 8^. Microvilli first appear on the apical surface of epithelial cells during the early stages of differentiation, and subsequently undergo rapid population expansion and packing to form mature brush borders ^9, 10^. Microvillus structure is highly stereotyped: each protrusion is supported by 30-40 actin filaments bundled in parallel such that their fast-growing barbed ends are oriented towards the distal tips ^6, 11^, which are marked by a complex containing epidermal growth factor pathway substrate 8 (EPS8) and insulin-receptor tyrosine kinase substrate (IRTKS) ^12, 13^. Parallel filament bundling is supported by fimbrin (also known as plastin-1), villin, espin, and mitotic spindle positioning protein (MISP) ^14–17^. Recent studies revealed that these factors occupy distinct regions along the length of the core bundle, suggesting a potential mechanism for regional tuning of physical properties ^17^. The core actin bundle is connected to the encapsulating membrane through membrane-cytoskeleton linkers, including ezrin and myosin-1a, which maintain membrane wrapping and increase long-term protrusion stability ^18, 19^. Despite significant progress toward defining the structure and composition of microvilli, mechanisms that control the growth of these protrusions remain poorly understood.

The morphologies of surface features such as microvilli are regulated by a balance of force-generating cytoskeletal elements and the force-resisting physical properties of the plasma membrane ^20, 21^. Endocytosis, the process by which cells internalize membrane and extracellular materials, is well positioned to impact this force balance as it involves localized membrane deformation that is potentiated by actin polymerization. The series of molecular events that drive actin assembly during clathrin-mediated endocytosis (CME) is well-studied in both yeast and mammalian cells ^22^. Synthesis of phosphatidylinositol 4,5-bisphosphate (PI(4,5)P_2_) at clathrin coated pits (CCPs) ^23^ triggers the recruitment of actin-binding proteins including neuronal-Wiscott-Aldrich syndrome protein (N-WASP)^24^ and actin related protein 2/3 complex (Arp2/3)^25^, capping protein ^26^, and actin-bundling ^27^ and severing proteins such as cofilin ^28^; these proteins work together to promote the polymerization and turnover of actin networks to drive the formation and scission of CCPs ^22^. Although CCPs can internalize without actin assembly, in high membrane tension scenarios such as the epithelial apical surface, actin polymerization counteracts the additional resistance to membrane bending ^29, 30^. Intriguingly, CCPs and microvilli share several molecular components such as actin-binding and adaptor molecules, suggesting a potential mechanistic link between these functionally and topologically distinct apical processes. For example, in solute transporting epithelial cells, PI(4,5)P_2_ is enriched in the apical membrane and has been shown to recruit and activate ezrin, a key membrane-cytoskeletal linker in the brush border ^31^. Actin-bundling proteins like fimbrin, which crosslink filaments in the microvillar core actin bundle, are also recruited to endocytic patches ^15, 27^. Mammalian actin binding protein 1 (mAbp1), with binding domains for both actin and clathrin, acts as a molecular bridge by physically linking the actin cytoskeleton to endocytic machinery ^32^. Notably, mAbp1 has also been shown to bind two molecules implicated in building microvilli: pacsin2 (also known as syndapin2) ^33^ and cordon bleu (COBL) ^34^. Previous studies established that knockout of pacsin2 in mice resulted in significantly shorter and fewer microvilli with reduced membrane wrapping on the surface of mature villus enterocytes ^35^. That work also revealed stalled endocytic intermediates, which accumulated between microvilli, suggesting a defect in endocytic activity. Although the mechanisms underlying these phenotypes remain unclear, when combined with the molecular links alluded to above, these studies collectively suggest that endocytic activity at the apical surface may be needed for normal microvillar growth.

Does CME control the growth of microvilli? To address this question, we used a combination of genetic and small molecule perturbations coupled with live confocal and ultrastructural imaging to examine microvillar dynamics and structure on the apical surface of brush border-building epithelial cell lines. Perturbation of CME in actively differentiating cells led to dramatic remodeling of microvilli; activation of CME depleted microvilli, whereas inhibition resulted in exaggerated protrusions containing more actin filaments and actin-binding proteins. Interestingly, volumetric live-cell imaging revealed that microvilli grow directly from CCPs, and scanning electron microscopy (SEM) also revealed a tight spatial association between apical surface pits and microvilli in both kidney- and intestine-derived transporting epithelial cells. Finally, inhibition of the Arp2/3 branched nucleation complex, which drives actin polymerization on coated pits, significantly reduced the accumulation of new microvilli on the surface of differentiating epithelial cells. Together, these data suggest that, across multiple epithelial models, CME, and its associated Arp2/3-based actin nucleation activity, controls the timing and location of microvillar growth as well as the dimensions of resulting protrusions. These findings redefine our understanding of microvillar biogenesis and, given the widespread appearance of CME in diverse biological systems, hold broad implications for understanding how cell surface morphologies are controlled.

## RESULTS

### Endocytic markers localize to the apical surface of actively differentiating epithelial cells

While previous studies established that the endocytic machinery localizes to the apical surface of functionally mature transporting epithelial cells, whether endocytosis plays a role in microvillar growth during epithelial differentiation remains unexplored ^36, 37^. We first turned to SEM to examine the surface of cells lining intestinal crypts, the site of microvillar growth in intestinal tissues, where nutrient transporting enterocytes comprise ∼95% of the epithelium. The intestinal crypt-villus axis offers a unique opportunity for imaging epithelial cells across all stages of differentiation; crypts house the transit amplifying cells (actively differentiating) while the villus is populated by mature epithelial cells (fully differentiated) ^38^. SEM of crypts in fractured samples of mouse small intestine revealed a gradual increase in microvillar density on the apical surface, suggesting that the brush border continuously packs with protrusions as cells migrate from the base of the crypt to the crypt-villus transition (Fig. 1A,B). We confirmed this point using immunofluorescent staining to probe for the microvillar-specific actin-bundler, villin, as a proxy for microvillar density. Consistent with the SEM data, confocal images revealed the apical villin signal increased nearly 2-fold from crypt base (magenta) to crypt-villus transition (green) (Fig. 1C,D), indicative of microvillar accumulation on the surface of these cells. We next stained for dynamin2 and pacsin2 to assess the abundance of these canonical endocytic markers in the crypts. The resulting confocal images revealed that, similar to villin, both markers demonstrated a gradual increase in signal, from the crypt base (magenta) to crypt-villus transition (Fig. 1C,D). Collectively, these data suggest that the accumulation of microvilli at the apex of actively differentiating cells spatially and temporally coincides with an increase in endocytic marker enrichment at the apical surface, well before cells are exposed to the contents of the lumen.

**Figure 1.**
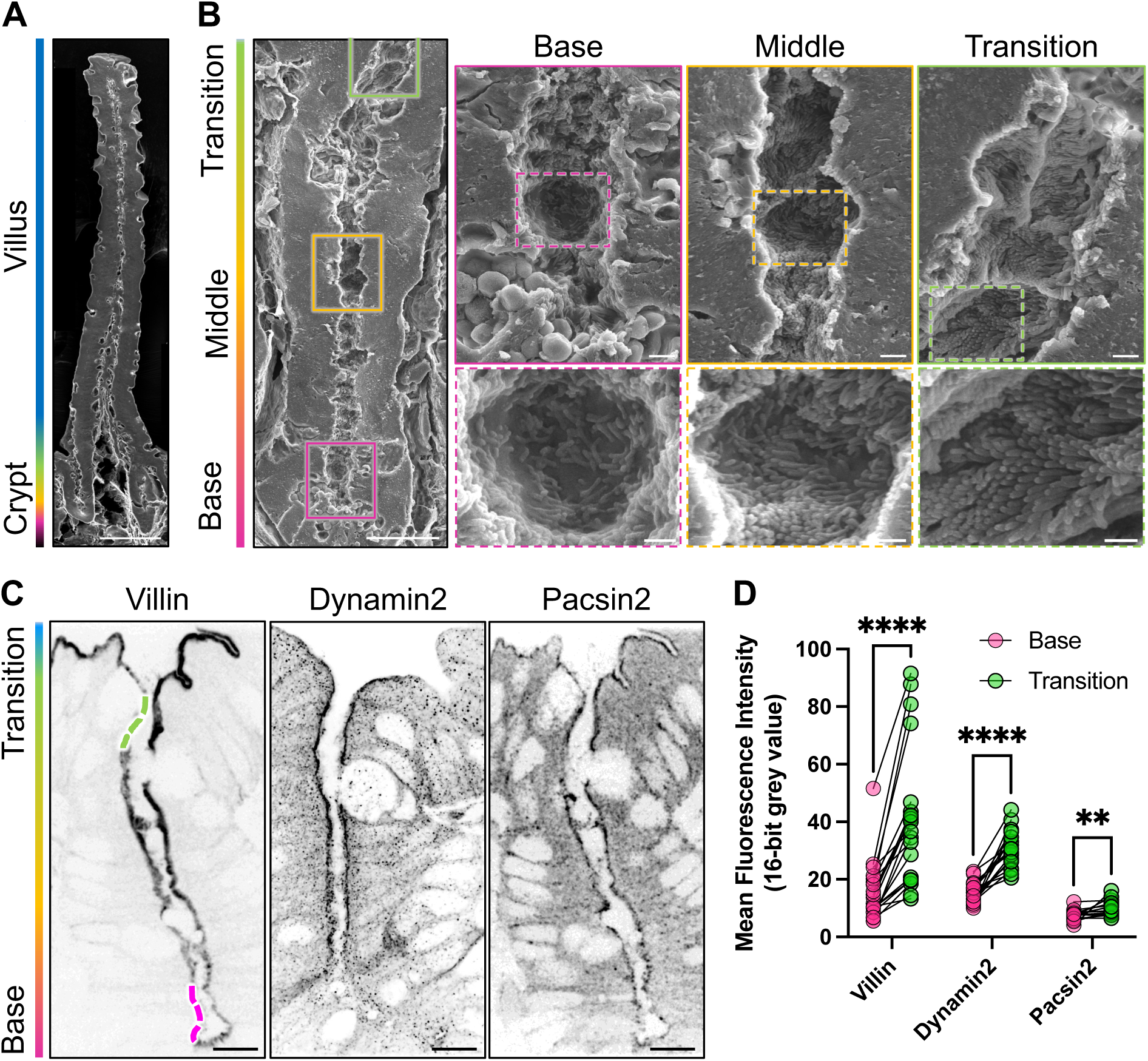
Endocytic proteins are apically localized in the crypts. (A) Scanning electron microscopy (SEM) of native mouse small intestine along full crypt-villus axis. Scale bar = 50 µm. (B) High magnification of the small intestinal crypt (scale bar = 10 µm) with zooms (base, middle, and transition (scale bars = 1 um (solid box zooms) and 500 nm (dashed box zooms)) showing the accumulation of microvilli along the transit amplifying zone. (C) Native mouse small intestine stained for villin, dynamin2, and pacsin2. Panels show single z-plane confocal image of intestinal crypts. Scale bar = 10 µm. (D) Mean brush border intensity measurements for each marker at the base of the crypt (magenta) and at the crypt-villus transition zone (green). n = 20-25 crypts per marker, 3 WT mice. Multiple Mann-Whitney tests; ****p = <0.0001, **p = 0.002.

### Induction of clathrin-mediated endocytosis depletes microvilli

Given the localization of clathrin-mediated endocytic components to the apical surface of differentiating epithelial cells in native intestinal crypts, we sought to determine if perturbing CME impacted microvillar growth. Here we employed the previously established “hook and anchor” tool to induce clathrin-mediated endocytosis ^39^. In this system, FKBP-tagged β2 subunit of AP-2 (β2) is recruited to an FRB-tagged CD8 membrane anchor upon addition of rapalog, resulting in robust formation and internalization of CCPs in a temporally controlled manner (Fig. 2A,B). We expressed this system in undifferentiated CL4 cells along with Halo-tagged espin, a microvillar actin bundling protein that specifically labels microvilli ^16^. Following endocytic induction, we performed live spinning disk confocal (SDC) imaging to monitor any effects on microvillar growth and structure (Fig. 2C). To measure the impact on core actin bundle structure, we generated linescans orthogonal to individual protrusions visualized in the Halo-espin channel, and calculated areas under the resulting intensity curves (AUC); the resulting metric is expected to reflect the diameter of the core bundle and the number of constituent actin filaments (Fig. 2D). Performing this analysis on cells before and 45 min after rapalog treatment revealed that inducing CME resulted in a ∼50% reduction in Halo-espin signal (Fig. 2E). Additionally, total microvillar surface coverage after endocytosis induction was also reduced relative to the GFP control (Fig. 2F). Together these data suggest that induction of CME negatively impacts microvillar growth and stability.

**Figure 2.**
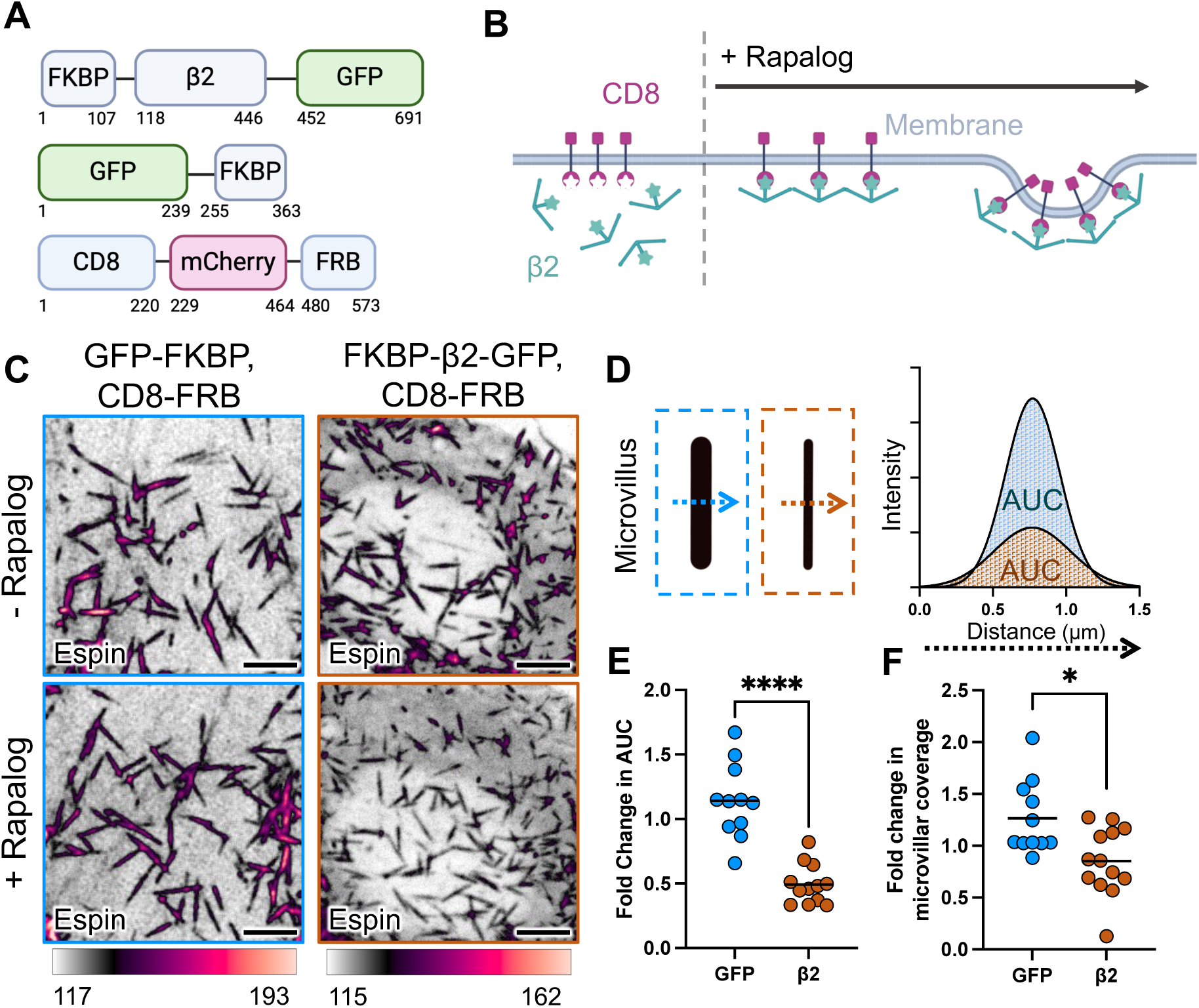
Induction of clathrin-mediated endocytosis depletes microvilli. (A) Cartoons depicting constructs used to induce clathrin-mediated endocytosis. Numbers represent amino acids. (B) Schematic depicting the rapalog-inducible system to dock the β2 subunit of AP-2 on the plasma membrane to induce endocytosis. (C) Confocal MIP of CL4 cell expressing Halo-espin (shown), CD8-FRB (not shown), and GFP-FKBP (not shown) or FKBP-β2-GFP (not shown) before and after 45 min rapalog incubation. Scale bar = 5 µm. (D) Schematic showing area under the curve (AUC) quantification. (E) Fold change in AUC of Halo-espin signal of microvilli post- vs. pre- rapalog induction. Area under the curve (AUC) was calculated as shown in D. 11-12 cells per condition taken from at least three experimental replicates, 120-150 microvilli per condition. Mann-Whitney test; ****p =<0.0001. (F) Fold change in area of microvillar coverage before and after 45 min rapalog incubation. N = 11-13 cells per condition, taken from at least three experimental replicates. Mann-Whitney test; *p=0.0218.

### Inhibition of clathrin-meditated endocytosis leads to the formation of exaggerated microvilli

We next sought to determine the impact of endocytic inhibition on microvillar assembly. For these studies, we used well established inhibitors to disrupt specific steps in the CME pathway and employed SDC microscopy to visualize the impact on protrusion growth and dynamics on the surface of CL4 cells expressing mCherry-espin^16, 40^. We inhibited CME by targeting both the initiation and scission steps of the pathway through chlorpromazine (CPZ) and dyngo, respectively. CPZ disrupts clathrin lattices on the plasma membrane and targets AP-2 to endosomes, while dyngo inhibits dynamin2, a GTPase critical for vesical scission during the final step of CCP invagination ^41, 42^ (Fig. 3A). CPZ treatment of undifferentiated CL4 cells resulted in dramatic exaggeration of microvillar structure (Movie S1); following 45 minutes of CPZ treatment, protrusions demonstrated a ∼1.5-fold increase in AUC measured from the espin signal and ∼2-fold increase in length (Fig. 3B-D, Movie S1). This effect was titrated across a range of CPZ concentrations in CL4 cells and resulted in a similar phenotype in CACO-2_BBE_ cells treated with CPZ (Fig. S1). In contrast, 45 minutes of dyngo treatment did not significantly impact microvillar morphology (Fig. 3B-D). To ensure that the observed phenotypic differences were not due to disparities in drug efficacy, we performed a transferrin uptake assay to assess CME activity following 45 minutes of treatment with either CPZ or dyngo. As expected, both endocytosis inhibitors significantly reduced transferrin uptake (Fig. S2). Thus, blocking endocytosis early in the pathway (Fig. 3A) promotes the growth of larger microvilli.

**Figure 3.**
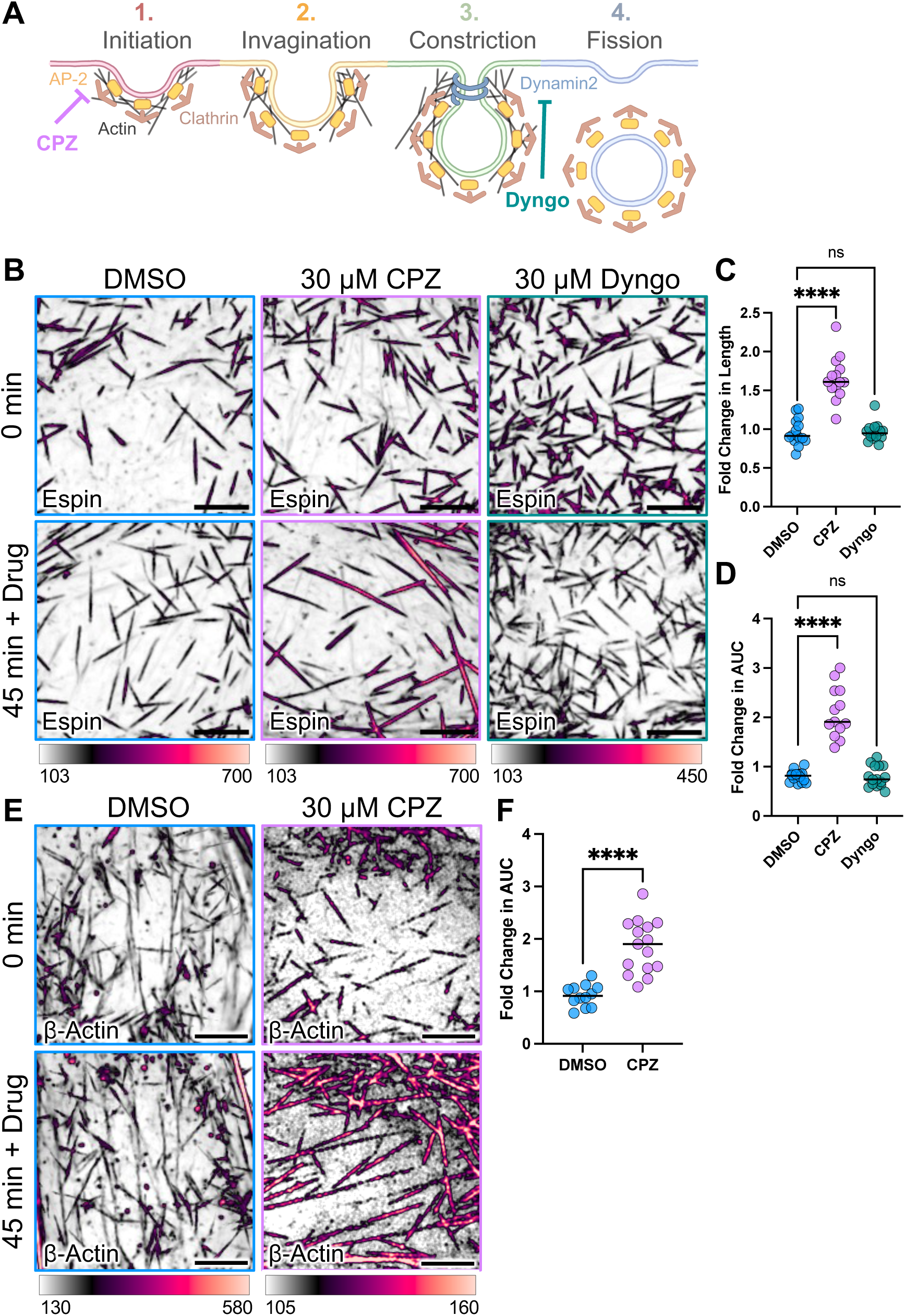
Inhibition of endocytosis results in exaggerated microvilli. (A) Schematic depicting chlorpromazine and dyngo mechanisms of action. (B) Confocal maximum intensity projection (MIP) of representative CL4 cells at 0 min and 45 min post DMSO (left), chlorpromazine (CPZ) (middle), and dyngo (right) treatment. Drug was added immediately after 0 min image was captured. Scale bar = 5 µm. (C) Fold change in length of microvilli post- vs. pre- drug treatments. Microvillar length was measured using mCherry-espin signal. n = 13-16 cells from at least three experimental replicates, 150-200 microvilli per condition. Kruskal-Wallis test with multiple comparisons; ****p = <0.0001. (D) Fold change in area under the curve of mCherry-espin signal of microvilli post- vs. pre- drug treatments. 14-16 cells from at least three experimental replicates, 150-200 microvilli per condition. Kruskal-Wallis test with multiple comparisons; ****p = <0.0001. (E) Confocal MIP of representative CL4 cells expressing mCherry-espin (not shown) and mNeonGreen-βActin at 0 min and 45 min post-DMSO or CPZ treatment. Scale bar = 5 µm. (F) Fold change in area under the curve of mNeonGreen-βActin signal of microvilli post- vs. pre- drug treatments. n = 15 cells from at least three experimental replicates, 150-200 microvilli per condition. Mann-Whitney test; ****p = <0.0001.

The increase in mCherry-espin signal in microvilli following CPZ treatment (Fig. 3B-D) suggests that growing core bundles incorporate more actin filaments in the absence of normal endocytic activity. To determine if this is the case, CL4 cells expressing mNeon-Green-ß-actin were treated with CPZ as described above (Fig. 3E). Consistent with the mCherry-espin readout, we observed a ∼2-fold increase in AUC, indicating that CPZ treatment indeed leads to higher levels of actin in exaggerated microvilli post-CPZ treatment (Fig. 3F). Labeling CPZ-treated cells with CellMask-DeepRed (CellMask), a plasma membrane dye, further revealed that exaggerated microvilli on these cells retained membrane wrapping, with the length of the wrapped region increasing with the longer core bundles formed under these conditions (Fig. 4A,B). We also examined the localization of ezrin, a major membrane-actin linker in these protrusions ^43^. AUC analysis of EGFP-ezrin signal in CPZ-treated cells further confirmed the increased diameter of exaggerated microvilli (Fig. 4C,D). Finally, we examined the impact of endocytic inhibition on two factors that reside at the distal tips of microvilli: EPS8 and IRTKS ^12^. Distal tip fluorescence intensity of EGFP-EPS8 or EGFP-IRTKS, quantified using Trackmate, revealed an enrichment of both EPS8 and IRTKS after CPZ treatment (Fig. 4E-H). Collectively, these data indicate that, under normal conditions, CME plays a role in limiting microvillar dimensions and actin content.

**Figure 4.**
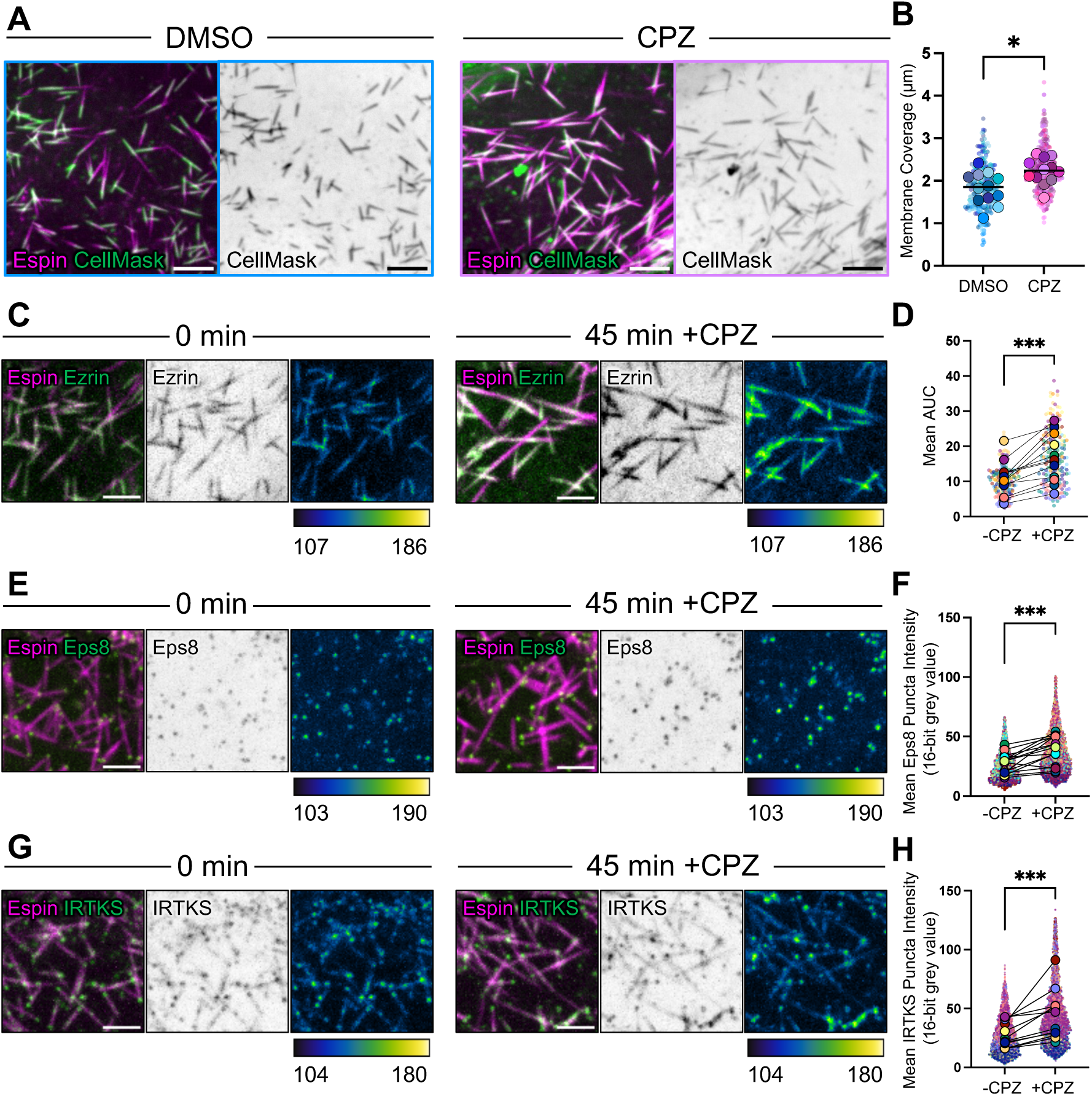
Exaggerated microvilli that assemble post-CPZ treatment retain canonical features. (A) Confocal maximum intensity projection (MIP) of CL4 cells expressing mCherry-espin stained with CellMask-DeepRed. Cells were first treated with either DMSO or CPZ, then incubated with CellMask for 15 minutes. Scale bar = 5 µm. (B) Superplot quantification of the length of membrane coverage along the microvillus after DMSO or CPZ treatment. Small, transparent points represent individual microvilli, larger opaque points show whole cell averages. (C, E, G) Confocal MIP of CL4 cells expressing mCherry-espin and (C) EGFP-ezrin, (E) EGFP-EPS8, and (E) EGFP-IRTKS at 0 min and 45 min. Single channel zooms show intensity-coded color profile. Intensity scales from low (dark blue) to high (yellow). Scale bar = 3 µm. (D) Quantification of ezrin AUC before and after CPZ treatment. N = 11 cells, 130-160 microvilli per condition. Mann Whitney U test; ***p =0.0002. (F,H) Superplot quantification of (F) Eps8 or (H) IRTKS intensity pre/post CPZ treatment. Small, transparent data points are individual puncta and larger, opaque data points are individual cells. N = 13-15 cells, 4000-6000 microvilli per condition. Mann Whitney U test; ***p = 0.0005.

### Exaggerated microvilli that form following CPZ treatment exhibit abnormal motility

Microvilli that extend from differentiating epithelial cells exhibit active motility across the apical surface, which is driven by the treadmilling of the supporting core actin bundles; this activity drives collisions between nascent protrusions, which in turn promote the formation of intermicrovillar adhesion complexes and microvilli packing ^44^. Given the increased length and filament content of microvilli following CPZ treatment, we sought to determine if these exaggerated protrusions exhibit aberrant motility. To address this question, we employed live SDC imaging to track the motility of individual microvilli on CL4 cells co-expressing mCherry-espin and EGFP-EPS8 (Fig. 5A). Mean square displacement analysis of 50 randomly selected microvillar trajectories per condition revealed parabolic curves consistent with a mixture of robust active motion and limited diffusion (Fig. 5D). Further, while we observed no significant changes in velocity nor track displacement after endocytosis inhibition, exaggerated microvilli did exhibit higher persistence of motion, which was reflected in the extended reach of individual trajectories when viewed in rose plots (Fig. 5B,C,E-G). Thus, inhibition of CME with CPZ leads to the growth of exaggerated microvilli that exhibit abnormal motility.

**Figure 5.**
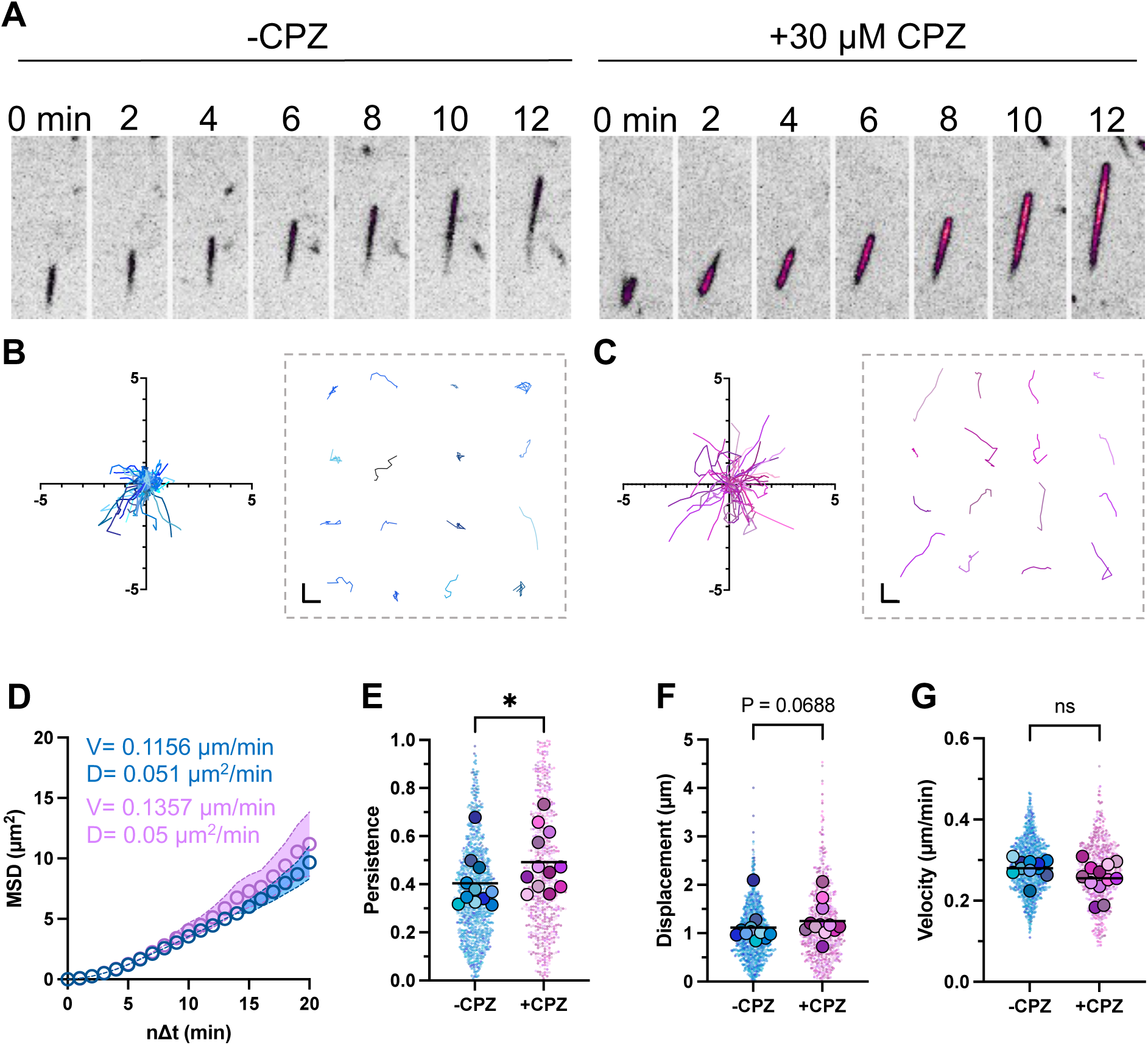
Exaggerated microvilli exhibit highly persistent motility. (A) Representative montage of microvillar motility and elongation before (left) and after (right) CPZ treatment. Single channels show intensity-coded color profile. Intensity scales from low (white) to high (yellow). Column width = 2 μm. (B) (left) Rose plot of 50 microvillar tracks over 10 min (μm units) in cells pre CPZ treatment. B. (right) 16 representative tracks isolated from B. Scale bar = 1 µm. (C) (left) Rose plot of 50 microvillar tracks over 10 min (μm units) in cells post CPZ treatment. C. (right) 16 representative tracks isolated from C. Scale bar = 1 µm. (D) Mean squared displacement analysis of 100 microvillar tracks imaged for 20 minutes every 1 minute. (E-G). Superplot quantification of persistence, displacement, and velocity, respectively in untreated and CPZ treated cell populations. Only tracks lasting exactly 10 minutes were analyzed. Small transparent points show individual microvilli, larger opaque points show per cell averages. n = 10-12 cells sampled from at least three independent experiments, 60-100 microvilli per cell. Mann Whitney test; P* = 0.0439.

### Global reduction of plasma membrane tension does not lead to the formation of exaggerated microvilli

Endocytic activity plays a global role in decreasing plasma membrane surface area and thus, increasing tension in this bilayer ^45^. Moreover, previous studies on CPZ suggest this compound intercalates into the plasma membrane, which is expected to reduce membrane tension ^46, 47^. To determine if global perturbations in membrane tension are responsible for the growth of exaggerated microvilli post-CPZ treatment, we treated CL4 cells with other compounds that are known to reduce membrane tension including 400 µM deoxycholate, 1% ethanol, and addition of 125 mM NaCl; all of these treatments decrease membrane tension as measured in tether force assays ^48–50^. However, none of these global perturbations induced the formation of microvilli with exaggerated dimensions (Fig. S3). These results suggest that endocytic events might instead locally regulate the growth and dimensions of microvilli.

### Microvilli grow from apical clathrin-coated pits

We next sought to determine if endocytic sites are spatially and temporally linked to microvillar growth events on the apical surface. To this end, we leveraged a recently developed SDC-based live imaging approach which allows for the visualization of individual microvillar growth events on the surface of sub-confluent CL4 cells ^9^. To screen for potential spatiotemporal links between microvillar growth and CME, we imaged mCherry-espin expressing CL4s co-expressing a range of EGFP-tagged endocytic components including pacsin2, clathrin light chain A (LCA), dynamin2, mAbp1, and PI(4,5)P_2_ (Fig. 6A,C,E and Fig. S4) ^32, 51–53^. To quantify the localization of endocytic markers at sites of microvillar growth, we analyzed the maximum intensity projected fluorescence signals of CME marker puncta that colocalized with elongating espin signals (representing the growth of new microvillar core bundles) over time. Strikingly, most endocytic markers we examined localized to the apical surface at future sites of microvillar growth ∼2-3 minutes prior to espin accumulation and elongation (Fig. 6B,D,F). One exception to this pattern was dynamin2, which briefly localized ∼1 minute after espin accumulation began, presumably marking the disassembly of the CCP (Fig. S4C). Following their localization to sites of microvillar growth, most endocytic components disassembled within a few minutes, a timeframe consistent with previously reported dynamics of CME ^54^. Together, these timelapse recordings suggest that the early steps of CCP formation create a specialized apical membrane domain that is permissive for microvillar growth.

**Figure 6.**
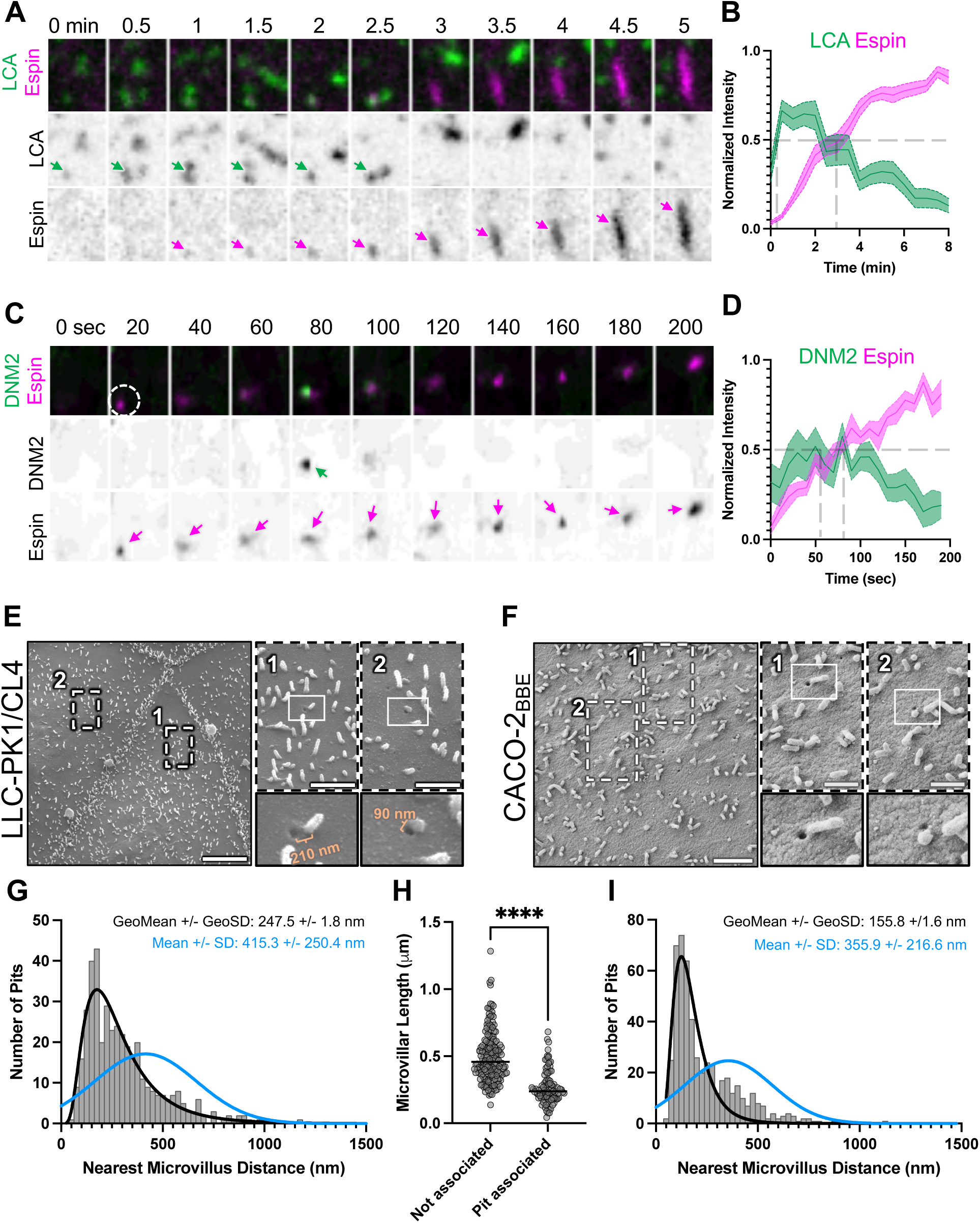
Microvilli grow from apical clathrin coated pits. (A) Montage of a single microvillus growth event in CL4 cells expressing EGFP-LCA and mCherry-Espin. Every column represents 30 seconds. Arrows mark the analyzed core bundle. Column width = 2.09 µm. (B) Average fluorescence intensity measurements over time. n = 21 growth events from 8 cells across 3 independent experiments. (C) Montage of a single microvillus growth event in CL4 cells expressing mCherry-Dynamin2 and GFP-Espin. Every column represents one minute. Arrows mark the analyzed core bundle. Column width = 2.09 µm. (D) Average fluorescence intensity measurements over time. n = 10 growth events from 4 cells across 3 independent experiments. (E) Scanning electron microscopy image of 0-DPC CL4 cells. Scale bar = 10 µm. Zooms 1 and 2 noted by bounding boxes. Zoom scale bar = 2 µm. (F) Scanning electron microscopy image of 0-DPC CACO-2_BBE_ cells. Scale bar = 2 µm. Zooms 1 and 2 noted by bounding box. Zoom scale bar = 1 µm. (G) Nearest neighbor distance measurements of from microvilli to closest endocytic pits on the surface of CL4 cells. Blue line denotes simulated nearest neighbor distances assuming random distribution of the same number of microvilli and endocytic pits across the same cell surface area. (H). Length of microvilli associated (<250 nm) vs not associated (>300 nm) with endocytic pits on the surface of CL4 cells. N = 175 microvilli in >300 category and 95 microvilli in <250 nm category. Mann Whitney test ****p = <0.0001. (I) Nearest neighbor distance measurements of from microvilli to closest endocytic pits on the surface of CACO-2_BBE_ cells. Blue line denotes simulated nearest neighbor distances assuming random distribution of same number of microvilli and endocytic pits across the same cell surface area.

### Ultrastructural imaging reveals juxtaposition of nascent microvilli and endocytic pits

To further examine the relationship between CCP formation and the growth of new microvilli, we employed SEM to visualize fine structural details of the apical surface of undifferentiated CL4 cells. Strikingly, these data showed that 51.1% of the pits that we could visualize were in proximity to microvilli (<250 nm) (Fig. 7A,C). To determine if the observed spacing was closer than expected based on randomly distributed pits and microvilli, we performed a nearest neighbor analysis, measuring from the center of every pit to the center of the base of the nearest microvillus. We also simulated cell surfaces and nearest neighbor measurements with the same number of pits and microvilli randomly distributed over the same area (Fig. S5). Remarkably, experimental nearest neighbor measurements exhibited a skewed distribution with a peak at ∼200 nm; in contrast, simulated random nearest neighbor measurements demonstrated a Gaussian distribution with a wider peak at ∼450 nm (Fig. 7C). Thus, the observed spacing between endocytic pits and microvilli cannot be explained by random co-occurrence and is instead consistent with two processes that are linked in space and time. We also noted that microvilli associated with endocytic pits were significantly shorter than microvilli further away, suggesting that core bundles mature as they as they disassociate from CCPs (Fig. 7D). Expanding the significance of this finding, we observed the same close juxtaposition of endocytic pits and microvilli on the surface of CACO2_BBE_ cells (67.1% of pits associated with microvillus) (Fig. 7B,E). These ultrastructural findings provide a compelling complement to our timelapse data, and strongly indicate that CME and microvillar biogenesis are spatially and temporally coordinated processes at the apical cell surface.

**Figure 7.**
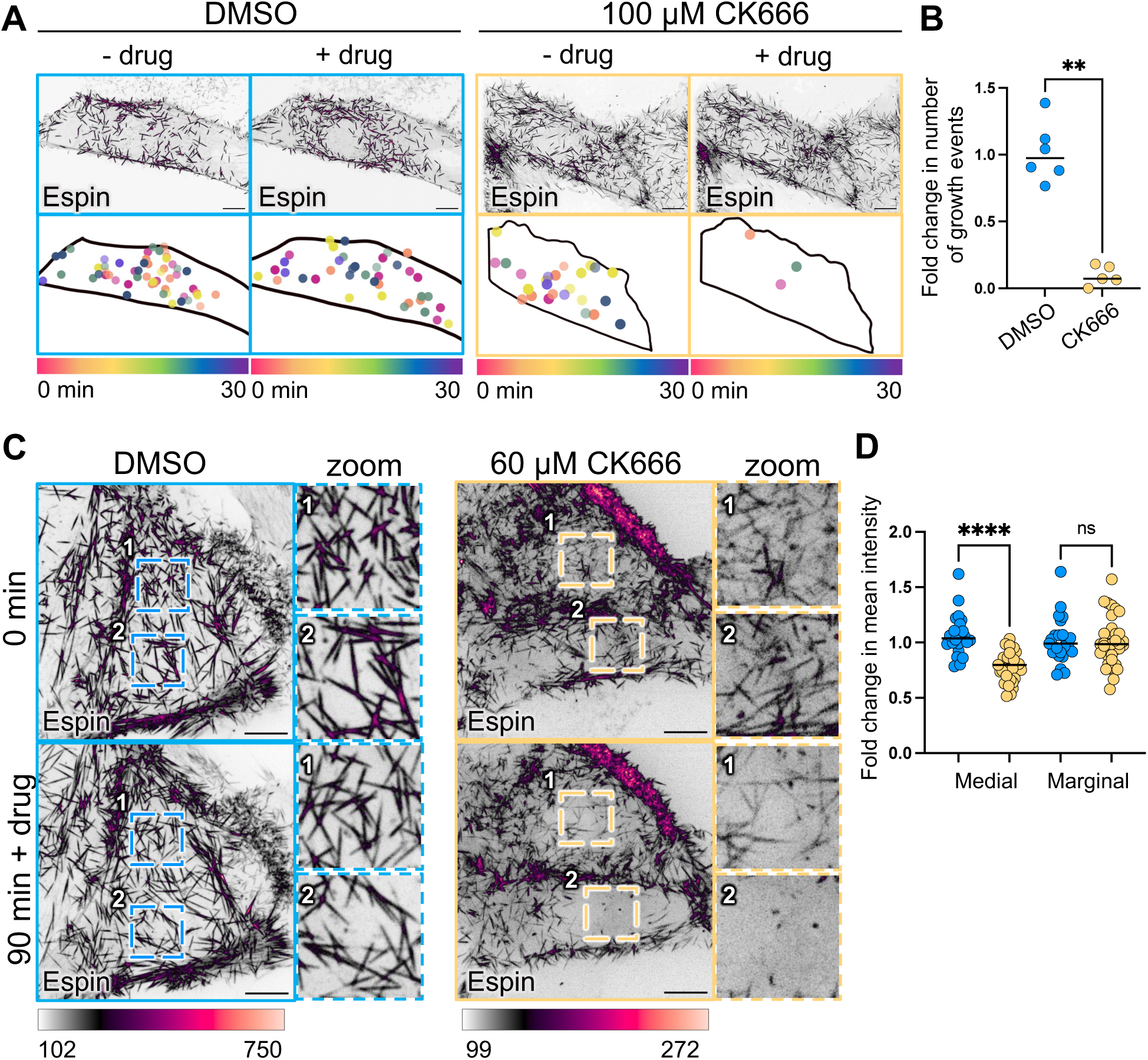
Arp2/3 complex inhibition decreases microvillar growth. (A) Confocal maximum intensity projection (MIP) of representative CL4 cells at 0 min and 45 min post DMSO (left) or 100 µM CK666. Cartoons beneath MIPs highlight growth events seen in Supplemental movie 2, color coded according to when growth event occurred during 30 minute imaging window. (B) Fold change in the rate of microvillar growth initiation. To define a fold change value for cells that had zero growth events following CK666 treatment, a value of 0.001 was added to all measurements. N= 5-6 cells per condition. Mann Whitney test **p = 0.0043. (C) Confocal MIPs of representative CL4 cells at 0 min and 90 min post DMSO (left) or 60 µM CK666 (right). Bounding boxes denote zoom insets. (D) Fold change in mean apical espin intensity of medial (middle) and marginal (junctional) microvilli. N = 23 cells in DMSO condition and 35 cells in CK666 condition. Ordinary one-way ANOVA with Šídák’s multiple comparisons test; ****p = <0.0001

**Figure 8.**
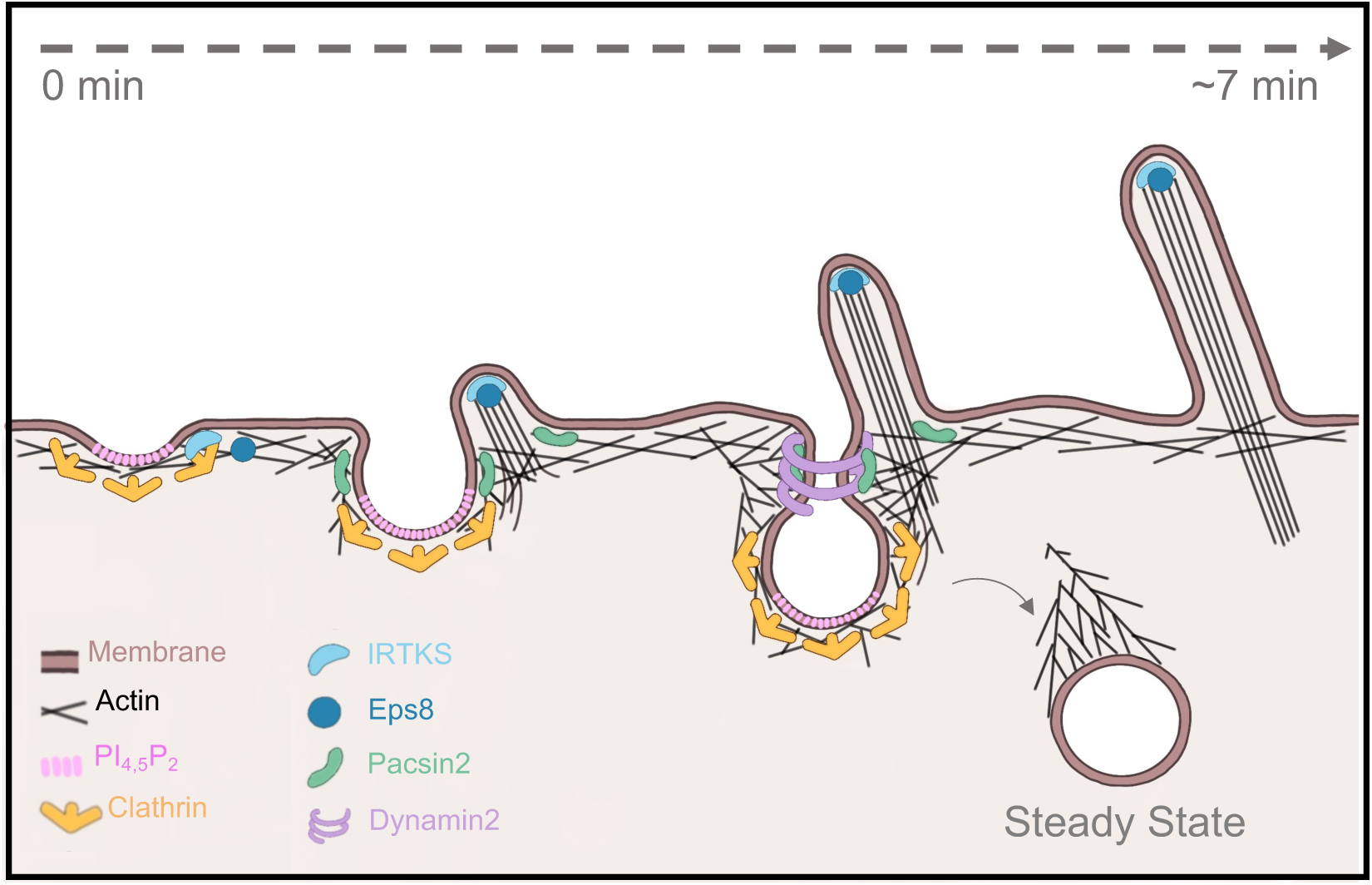
Schematic depicting proposed model of clathrin mediated endocytosis and microvillar biogenesis sharing materials during epithelial cell differentiation.

### Arp2/3 complex inhibition decreases the growth of new microvilli

Actin filament assembly on the surface of CCPs, driven by the Arp2/3 branched nucleation complex, is critical for promoting pit closure and internalization under conditions of high membrane tension, like that experienced at the apical surface ^29, 30^. Given the close association of nascent microvilli and endocytic pits revealed in our confocal and SEM imaging studies, we wondered if Arp2/3-generated actin filaments on the surface of CCPs directly support the formation of microvillar core actin bundles. Because the actin nucleators that support microvillar growth have yet to be identified, we were eager to test this idea. Toward this end, we used SDC-based live imaging to visualize individual microvillar growth events on the surface of sub-confluent CL4 cells ^9^ in the presence of CK-666, a potent inhibitor of Arp2/3 nucleation activity ^55^. Time-lapse acquisitions over the course of 30 min revealed a 10-fold reduction in the rate of microvillar growth initiation following CK-666 treatment (Fig. 7A,B). When acquisitions were run for longer periods of time (90 min), analysis of total microvillar content on the apical surface also revealed a striking decrease in protrusions, suggesting that the reduced growth rates following CK-666 treatment also led to reduced accumulation of protrusions long-term (Fig. 7C,D). Together, these results suggest that actin filaments generated by CCP-associated Arp2/3 are remodeled to form the parallel, polarized core bundles that support microvilli.

## DISCUSSION

Despite the extensive characterization of microvillus architecture and composition, our understanding of how cells control the growth and resulting dimensions of these protrusions remains limited. Here, we establish apical endocytic activity as a critical regulator of the location, timing, and dimensions of nascent microvilli. Given the overlapping molecular machinery and the role of the actin cytoskeleton and membrane bending in both apical CME and microvilli, we sought to test the hypothesis that microvillar growth is linked to apical endocytic activity. Indeed, our analysis of endocytic markers (pacsin2 and dynamin2) in native intestinal crypts revealed that differentiating cells experiencing rapid microvillar growth also exhibit robust apical localization of endocytic machinery. This finding revealed that apical endocytic activity is well positioned to influence microvillar assembly early in the time course of epithelial differentiation.

How might CME impact microvillar growth on the apical surface of differentiating epithelial cells? Previous studies collectively indicated that experimental depletion of microvilli is difficult, underscoring the robustness of this evolutionarily ancient process ^56–59^. However, we found that inducing high levels of CME through the recruitment of the β2 subunit of AP2 to the plasma membrane led to a striking depletion of microvillar actin content and coverage across the cell surface. Our results echo those from previous studies which showed that overexpression of clathrin heavy chain perturbed the morphology of microvilli on the surface of Madin-Darby Canine Kidney (MDCK) cells, as assessed using transmission electron microscopy ^30^. A superficial interpretation of these findings is that apical CME influences microvillar growth by impacting apical membrane tension. Yet, in the current study, global perturbations of membrane tension using orthogonal methods failed to disrupt microvillar architecture. An appealing alternative model is that apical CCPs and microvillar assembly mechanisms share resources, such as specific protein machinery, membrane lipids, or actin filaments, and over- or underutilization of these resources by CME compromises the growth of protrusions from the apical surface. We elaborate more on the experimental support for this idea below.

If CCPs and microvilli share resources during their assembly, then inhibition of CME would be expected to divert those components into the microvillar growth pathway, and potentially result in more protrusions and/or larger structures. Indeed, when we inhibited CME using CPZ, a compound that inhibits clathrin recycling to the plasma membrane, new microvilli grew with exaggerated dimensions, appearing longer and with increased diameter, suggesting they contained more actin filaments. Protrusions assembled following CPZ treatment retained key structural features of canonical microvilli, including ezrin-mediated membrane wrapping, and distal tip localization of EPS8 and IRTKS. Interestingly, another endocytic inhibitor, dyngo, did not have the same effect, although this may be explained by the distinct mechanisms by which CPZ and dyngo block CME (Fig. 3A)^41, 42, 60^. CPZ prevents the formation of coated pits by disrupting AP-2 recruitment to the plasma membrane. In contrast, dyngo treatment allows for pit formation and leads to stabilization of pits on the plasma membrane by inhibiting vesicle scission. The stalled endocytic intermediates created by dyngo still sequester resources such as actin filaments, as suggested in previous studies ^61, 62^, potentially explaining why this treatment does not lead to microvillar overgrowth.

The apical surface phenotypes associated with gain and loss of CME function – depletion and exaggeration of microvilli, respectively - indicate that CCP formation and microvillar growth are mechanistically linked, potentially through a shared requirement for a molecular resource such as actin. This link could operate globally with these pathways engaging in long-range competition for the same pool of actin monomers, as suggested in previous studies on actin allocation ^63^. Alternatively, this link could operate locally, with both pathways sharing resources at the same time *and* in the same location at the apical membrane. This latter possibility is supported by our live imaging studies of epithelial cells early in differentiation. Using SDC microscopy, we defined a time course for the recruitment of endocytic machinery, which appeared in diffraction limited puncta on the apical surface in events that lasted several minutes as expected^54, 64^. Remarkably, these puncta also marked future sites of microvillar growth, with some CME markers arriving 2-3 minutes in advance of growth initiation. All CME markers examined in this assay demonstrated significant temporal and spatial overlap with the microvillar core bundle marker, espin, suggesting that resource sharing could occur at discrete sites that are scaled at or below the diffraction limit (< 200 nm). Interestingly, we noted that not all CCPs progressed to form microvilli. Given the variation in composition of CCPs, it may be that only a subset of pits are equipped to support microvillar growth^65–67^. What distinguishes these pits from their counterparts? One intriguing possibility lies in the mechanism of pit closure. Studies have identified three distinct closure types for CME: symmetric, asymmetric, and undefined, with asymmetric being the most prevalent (accounting for roughly 70% of pits) ^62, 68^. During asymmetric closure, membrane bulges outward on one side of the CCP propelled by actin polymerization, progressively covering and closing the pit. This contrasts with symmetric closure (∼20% of events), where the pit slowly closes in an isotropic manner^62, 68^. The specific mode of pit closure and/or the presence of asymmetric F-actin accumulation might dictate whether that pit ultimately leads to microvillus assembly^29, 62^.

We also surveyed the apical surfaces of fixed epithelial cell monolayers using SEM, with the goal of examining large numbers of endocytic pits and characterizing their spatial relationship with microvilli. Analysis of undifferentiated CL4 and CACO-2_BBE_ monolayers revealed that >50% of pits were within 250 nm of a microvillus (CL4: 51.1%, CACO-2 _BBE_: 67.1%), a distance comparable to the diameter of the diffraction limited puncta that we observed in our live confocal imaging studies. Mathematical simulations revealed that such close juxtaposition of endocytic pits and microvilli could not be explained by the random positioning of these objects across the apical surface. We also found that pit-associated microvilli were significantly shorter relative to more distal protrusions. This observation could indicate that pit-associated microvilli represent younger intermediates, whereas longer and more distal microvilli represent fully formed protrusions at steady-state length. In combination with the CME gain and loss of function studies and the timelapse observations described above, these data lead us to propose a model whereby microvilli form at sites that are immediately adjacent (within 250 nm) to pre-existing CCPs (Fig. 6F).

Why do CCPs serve as preferential sites for microvillar assembly? From several perspectives, the local biochemical and physical landscape generated during endocytic pit formation appears well-suited for supporting protrusion growth. For example, CCPs contain a significant F-actin coat generated by the Arp2/3 branched nucleation complex, and the protein machinery that drives microvillar growth could harness those polymers, elongating and bundling them to form core actin bundles. Interestingly, molecular counting studies revealed that ∼200 Arp2/3 complexes localize to CCPs, and simulations suggest that this population would be sufficient to generate ∼150 filaments for driving pit internalization^69^. Based on ultrastructural studies of the enterocyte brush border, we know that microvillar core actin bundles contain only ∼30-40 filaments ^11^, so the scale of the CCP-associated actin filament network is large enough to support the growth of a few microvilli. Inhibition of the Arp2/3 branched nucleation complex in CL4 cells significantly reduced the growth and accumulation of new microvilli on the surface of differentiating epithelial cells, suggesting that the actin filaments in the microvillus core bundle are derived from Arp2/3 generated networks (Fig. 7). The general concept of co-opting filaments from an existing network to form a new structure with distinct architecture is similar to the ‘convergent elongation’ model that was originally proposed by Svitkina and colleagues to explain the formation of filopodia on the surface of crawling cells^70^. In that scenario, the actin filaments that support filopodia originate deep in the lamellipodial meshwork, which is generated by Arp2/3 activity. During filopodial growth, a small number of lamellipodial filaments are protected from capping by barbed end elongation factors (Ena/VASP, mDia) and therefore allowed to elongate and become crosslinked by fascin to create a mechanically robust core bundle^70^. In the context of microvillar growth, EPS8 and/or IRTKS might function as elongation factors^12^, whereas espin and other factors (MISP, fimbrin, and villin) could participate in filament bundling^14–17^. Whether a similar mechanism is operational at the smaller scale of CCPs and leads to formation of microvilli will require deeper ultrastructural investigation.

SEM images revealed that when an endocytic pit is adjacent to a microvillus, the stoichiometry is almost always 1:1, i.e. one microvillus per pit. Thus, in addition to the availability of actin filaments, other constraints probably limit the number of new microvilli that form during these events. In our studies, we also observed enrichment of PI(4,5)P_2_ before microvilli begin to grow, suggesting a potential role for specific lipids in microvillar biogenesis. The local accumulation of specific lipids or membrane-associated proteins is known to impact membrane curvature and tension^71^, and these physical factors likely impact microvillar growth. Future investigations taking advantage of next generation membrane tension probes might explore this possibility.

The link between CME and microvillar growth revealed in this work points to a fascinating evolutionary adaptation in epithelial biology. While CME is a ubiquitous cellular process shared by all eukaryotes, microvilli are specialized structures that emerged later in evolution, with their earliest known appearance in choanoflagellates^3, 72^. The evolution of endocytosis provided a crucial foundation for membrane maintenance, cell signaling, and nutrient uptake^73, 74^. We propose that, over time, this actin-dependent process was co-opted to support the formation of actin-based protrusions such as microvilli. If true, this would represent an elegant example of evolutionary exaptation, where existing processes are repurposed to meet new functional demands. The model we propose here might also offer an explanation for growth of more elaborate microvilli-like structures, including the stereocilia that extend from the apical surface of mechanosensory hair cells found in the cochlear and vestibular systems.

## ACKNOWLEDGEMENTS

The authors thank all members of the Tyska laboratory for their constructive feedback. Microscopy was performed in part by the Vanderbilt Cell Imaging Shared Resource. This study was supported by the NIH grants R01 DK095811 (M.J.T.), R01 DK125546 (M.J.T.), and R01 DK111949 (M.J.T.).

## METHODS

### RESOURCE AVAILABILITY

#### Lead contact

Further information and requests for resources and reagents should be directed to the lead contact, Matthew J. Tyska.

#### Materials availability

Plasmids generated in this study will be made available from the lead contact on request.

#### Data code availability

No large-scale datasets or new code were generated in this study.

### EXPERIMENTAL MODEL AND SUBJECT DETAILS

#### Cell culture models

LLC-PK1-CL4 (porcine kidney proximal tubule) cells were grown in 1X high glucose DMEM containing 2mM L-glutamine (Corning #10-0130CV) supplemented with 10% fetal bovine serum (FBS) (R&D Systems) and 1% L-Glutamine (Corning # 25-005-Cl). CACO-2_BBE_ cells were grown in 1x high glucose DMEM containing 2mm L-glutamine supplemented with 20% FBS. Cells were maintained in treated plastic culture flasks incubated at 37 °C and 5% CO_2_. Cells were tested for mycoplasma monthly using the MycoStrip Mycoplasma Detection Kit (Invivogen #rep-mys-50).

### METHOD DETAILS

#### Cloning and Constructs

All overexpression constructs listed in this paper were previously generated and/or obtained as noted in the key resources table.

#### Cell line generation

LLC-PK1-CL4 cells stably expressing pmCherry-espin were previously generated using the stable cell line generation protocol as described in Gaeta et al., 2022. Briefly, cells were transfected with Lipofectamine 2000 following manufacturer protocol in a T25 cell culture flask. The next day, cells were split up to a T75 flask and treated with 1 mg/mL G418 sulfate for antibiotic selection. Cells were maintained in culture under constant G418 selection to create the stably expressing cell line.

#### Transient transfection

LLC-PK1-CL4 cells were seeded in 6 well plates and transfected at ∼90% confluency using Lipofectamine2000 with 3 µg DNA. 4 hrs after incubation with the lipid complex, cells were replated on CellVis 35mm 1.5# glass bottom dishes for imaging the following day.

#### Immunofluorescence staining

*Tissue:* Tissue sections were deparaffinized and rehydrated through a descending ethanol series and subjected to antigen retrieval in a heated buffer at pH 9. Following cooling, sections were washed and blocked with 10% normal goat serum (NGS) and probed for villin, pacsin2, or dynamin2 in 1% NGS overnight at 4 °C. The following day, sections were washed, incubated at room temperature with secondary antibodies, dehydrated through an ascending ethanol series, and mounted in ProLong Gold mounting medium.

#### Live imaging microscopy

Nikon Ti2 inverted light microscope equipped with a Yokogawa CSU-X1 spinning disk head, a Photometrics Prime 95B sCMOS camera, and 488 nm, 561 nm, and 647 nm excitation lasers. A 60x/1.49 NA or 100x/1.49 NA TIRF oil immersion objective was used for all experiments. Cells were maintained in a stage top incubator at 37 °C with 5% CO2 (Tokai Hit). Z stacks of 5-8 µm using a 0.2-0.4 µm step size were collected using a triggered NIDAQ piezo Z stage.

#### Scanning electron microscopy (SEM)

##### Tissue

5 mm murine duodenal sections were fixed in 2.5% glutaraldehyde and 4% paraformaldehyde in SEM, washed, and then embedded in OCT. To ensure stability of the explant lumens, samples were gently passed through 3 rounds of fresh OCT. Samples were then frozen in cryomolds and stored at -80 °C. Frozen explants were sectioned, melted onto stainless steel AFM discs, and processed through OsO4, ddH2O, and subsequently dehydrated through graded ethanol series. Detached sections were recovered and adhered to aluminum SEM specimen stubs. SEM imaging was performed on a Quanta 250 environmental SEM or Zeiss Crossbeam 550. For more detailed methods, see Meenderink et al^44^.

##### Cells

Caco-2_BBE_ cells were plated at 70% confluence on Thermanox coverslips in a 6-well dish. 24 hrs post-plating, cells were washed 3X with PBS to remove all media and were fixed in a buffer of 2.5% glutaraldehyde and 0.1 M CaCl_2_ free cacodylate overnight at room temperature. Coverslips were washed 3X in 0.1 M CaCl_2_ free cacodylate buffer and incubated in pre-warmed 1% tannic acid in cacodylate. Following washes with 0.1 M cacodylate, coverslips were incubated for 1 hour in 1% OsO_4_. After washes with ddH_2_O, they were incubated for 30 min with 1% uranyl acetate in ddH_2_O. Finally, coverslips were washed with ddH_2_O 3X for 5 min to remove uranyl acetate before drying with a series of graded EtOH washes. After samples were dried using critical point drying and vented for 20 min, coverslips were mounted on SEM specimen stubs via a conductive adhesive tab and sputter coated prior to imaging. SEM imaging was performed on a Zeiss Crossbeam 550 with a voltage of 2 keV and an SE2 detector. All reagents were purchased from Electron Microscopy Sciences.

#### Transferrin uptake assay

LLC-PK1-CL4 cells expressing mCherry-espin were seeded on acid-washed #1.5 coverslips at ∼60% confluence in a cell culture 6-well treated plate on day 0. The next day, cells were treated with 30 µM CPZ, 30 µM Dyngo, or DMSO and incubated for 45 min. At the 45-min mark, transferrin conjugate was added to the cells at 25 µg/mL and incubated for 20 min. After 20 min, the cells were fixed in 4% paraformaldehyde diluted in PBS and incubated at 37 °C for 15 min. Coverslips were subsequently mounted on slides in Prolong Gold antifade reagent and imaged using spinning disk microscopy.

### QUANTIFICATION AND STATISTICAL ANALYSIS

#### Statistical testing

All statistical testing was performed in GraphPad Prism 9 by running a Mann Whitney test or a paired t-test for pairwise comparisons or Kruskal-Wallis to compare more than two conditions. For all figures, exact n values and their definitions are reported in the figure legends.

#### Time series analysis of microvillus growth

Microvillar growth was analyzed as previously described^9^. Briefly, a bounding ROI was manually drawn around the signal of interest and mean intensity was measured for each timeframe. Mean intensity values were normalized, with the smallest value equal to 0 and the largest equal to one. Normalized values were then averaged and plotted with SEM.

#### Area under the curve analysis

To measure the impact on core actin bundle structure, we generated linescans (5 pixels in width, 1.54 µm in length) orthogonal to individual protrusions visualized in the Halo-EPSN channel. Linescans were drawn to ensure lines only crossed through one protrusion per line. Areas under the resulting intensity curves (AUC) were then calculated in prism, averaged and plotted; the resulting metric is expected to reflect the diameter of the core bundle.

#### EGFP-EPS8 and EGFP-IRTKS puncta intensities

Mean puncta intensities were measured using Nikon Elements’ “Spot Detection” function.

#### Microvilli tracking using EGFP-EPS8 puncta

Raw 3D movies were converted into maximum intensity projections, and the EPS8 channel (488) was isolated. The isolated channel movie was loaded into Trackmate and puncta were identified using a DoG detector with an estimated object diameter of 500 nm. Identified spots were thresholded based on background intensity and track length. Tracks beginning on the first frame or ending on the last frame were excluded. Track and spot data were then exported to Excel and analyzed and plotted in Prism as radial XY positions over time by subtracting each position in X or Y from their respective point position at time 0, making the first position (0,0). Persistence, track displacement, and linear velocity were taken from tracks lasting exactly 10 min.

#### Mean squared displacement analysis

EPS8 puncta trajectories as a proxy for microvillar motion were analyzed using mean squared displacement (MSD) analysis as previously described^44^. MSD curves were subsequently fit with an active motion model in the form MSD(nDt) = 4Dt + V^2^* nDt^2^, where D is the diffusion coefficient and V is the velocity of active motion ^75^. Curve fitting and sum-of-squares F-tests were performed using PRISM v.9.

#### Nearest neighbor expected distribution simulation

To determine the expected nearest neighbor distance between pits and microvilli, a random population of microvilli and endocytic pits were simulated within a rectangular area using a custom python script. The number of objects in each population was set to match the observed experimental densities. Euclidean distances between each endocytic pit and every microvillus were measured and the subsequent minimum distance (nearest neighbor) for each endocytic pit was identified. Each simulation (CACO-2_BBE_ and CL4) was run 10 times and then averaged (Extended Data Fig. 5).

## KEY RESOURCES TABLE

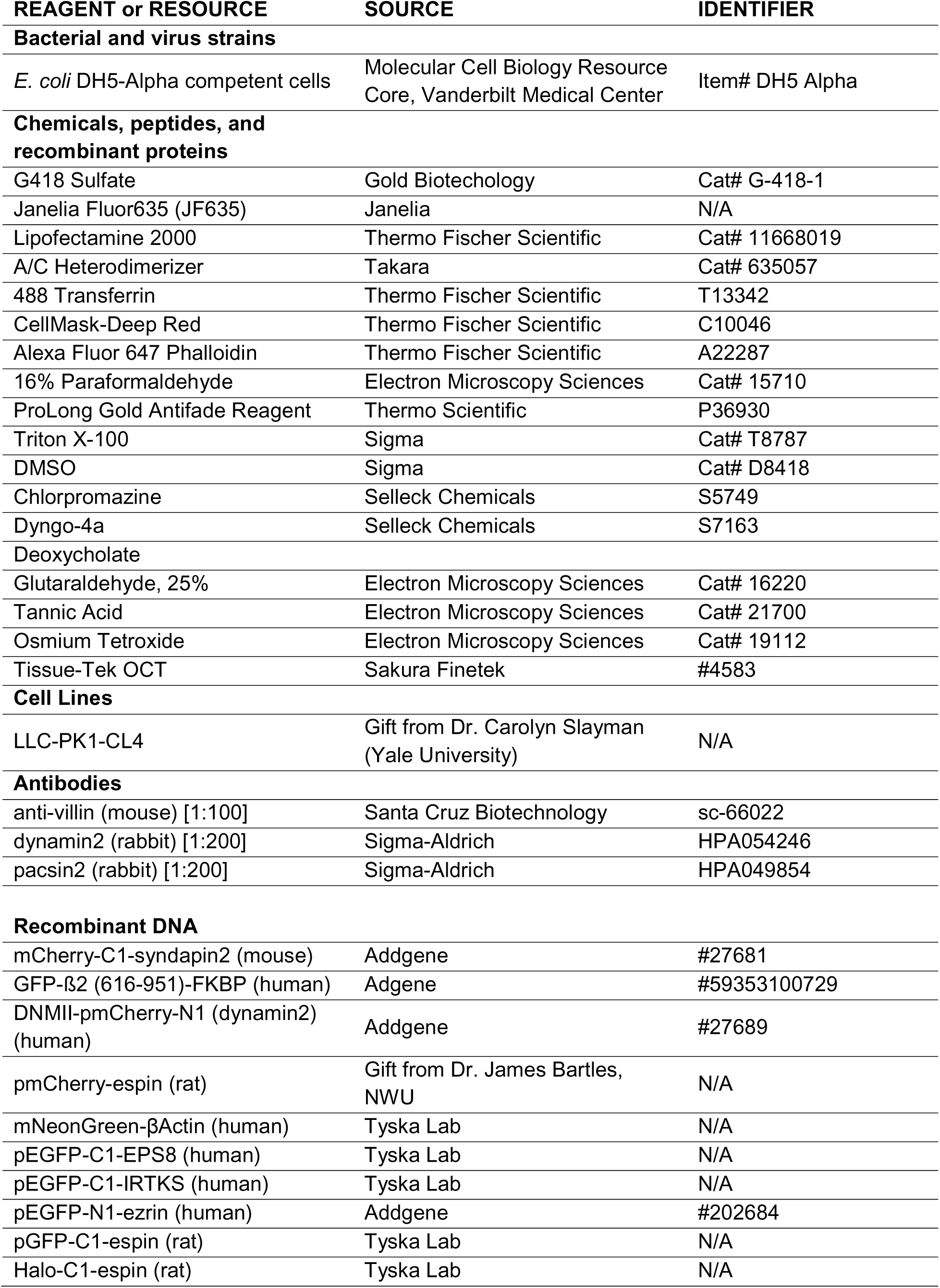
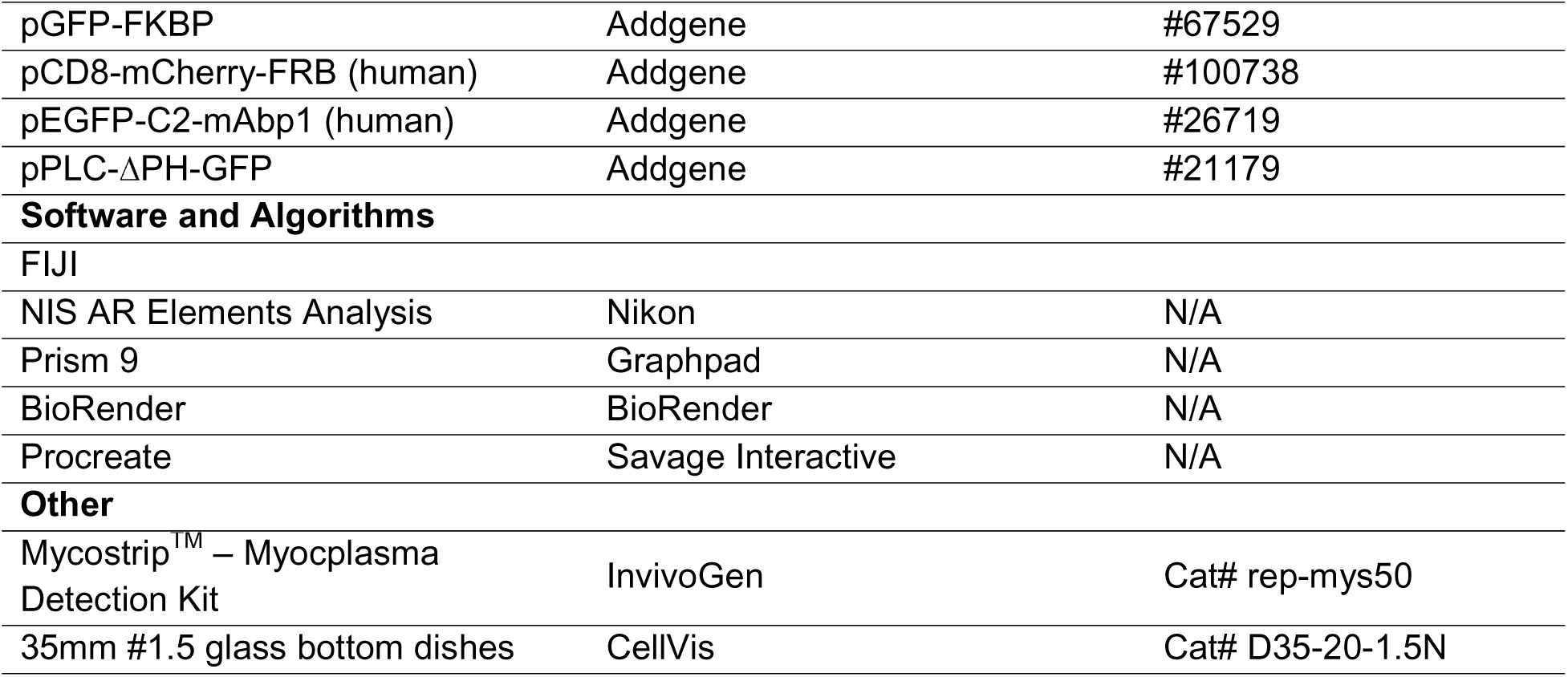

## SUPPLEMENTAL DATA

**Movie S1, related to Figure 3**. Inhibition of endocytosis results in exaggerated microvilli. Live cell movie of maximum intensity projection (MIP) confocal time series of 3 CL4 cells expressing mCherry-espin. CPZ was added at T = 30 min. Time hour: min: sec. Single channels show intensity-coded color profile. Intensity scales from low (white) to high (yellow). Scale bar = 5 µm.

**Movie S2, related to Figure 7. Arp2/3 complex is required for microvillar growth.** Live cell movie of maximum intensity projection (MIP) of confocal time series of 2 CL4 cells expressing mCherry-Espin. DMSO (left) or 100 µM CK666 (right) was added at T = 30 min. Cyan spots which accumulate highlight microvillar growth events. Images were captured at 1 frame per 1 minute. Time min: sec. Single channels show intensity-coded color profile. Intensity scales from low (white) to high (yellow).

**Figure S1.**
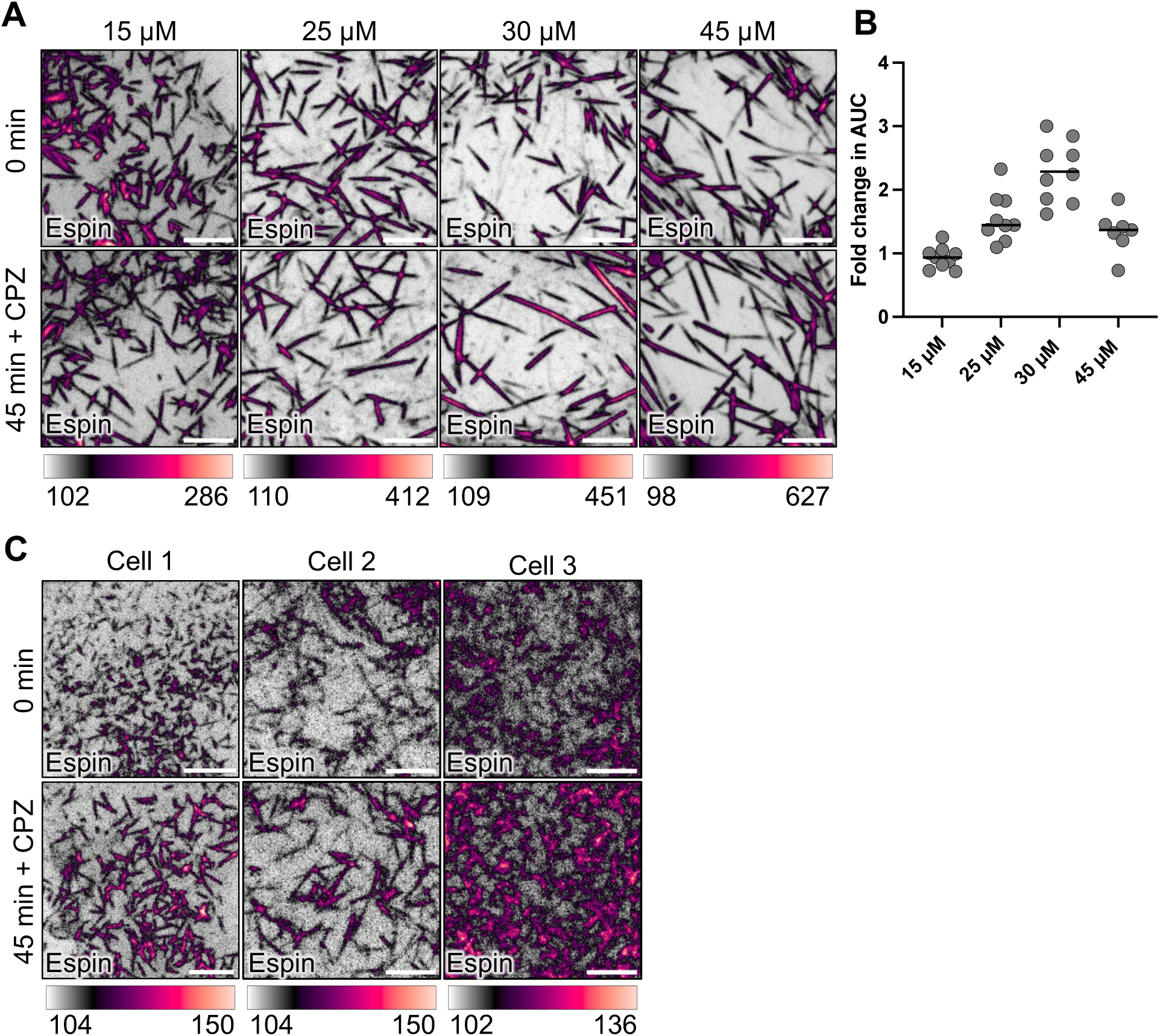
CPZ treatment phenotype is titratable and robust across model systems. (A) Confocal maximum intensity projection (MIP) of representative CL4 cells at 0 min and 45 min after varying concentrations of CPZ treatment. Drug was added immediately after 0 min image was captured. Single channels show intensity-coded color profile. Intensity scales from low (white) to high (yellow). Scale bar = 5 µm. (B) Fold change in area under the curve of mCherry-espin signal of microvilli post- vs. pre- CPZ treatment. 8-10 cells from at least three experimental replicates, 100-150 microvilli per condition. Images and data for the 30 µM CPZ treatment were taken from figure 3. (C) MIPs of 3 representative CACO_2_-BBE cells at 0 min and 45 min after treatment with 30 µM CPZ. Drug was added immediately after 0 min image was captured. Single channels show intensity-coded color profile. Intensity scales from low (white) to high (yellow). Scale bar = 5 µm.

**Figure S2.**
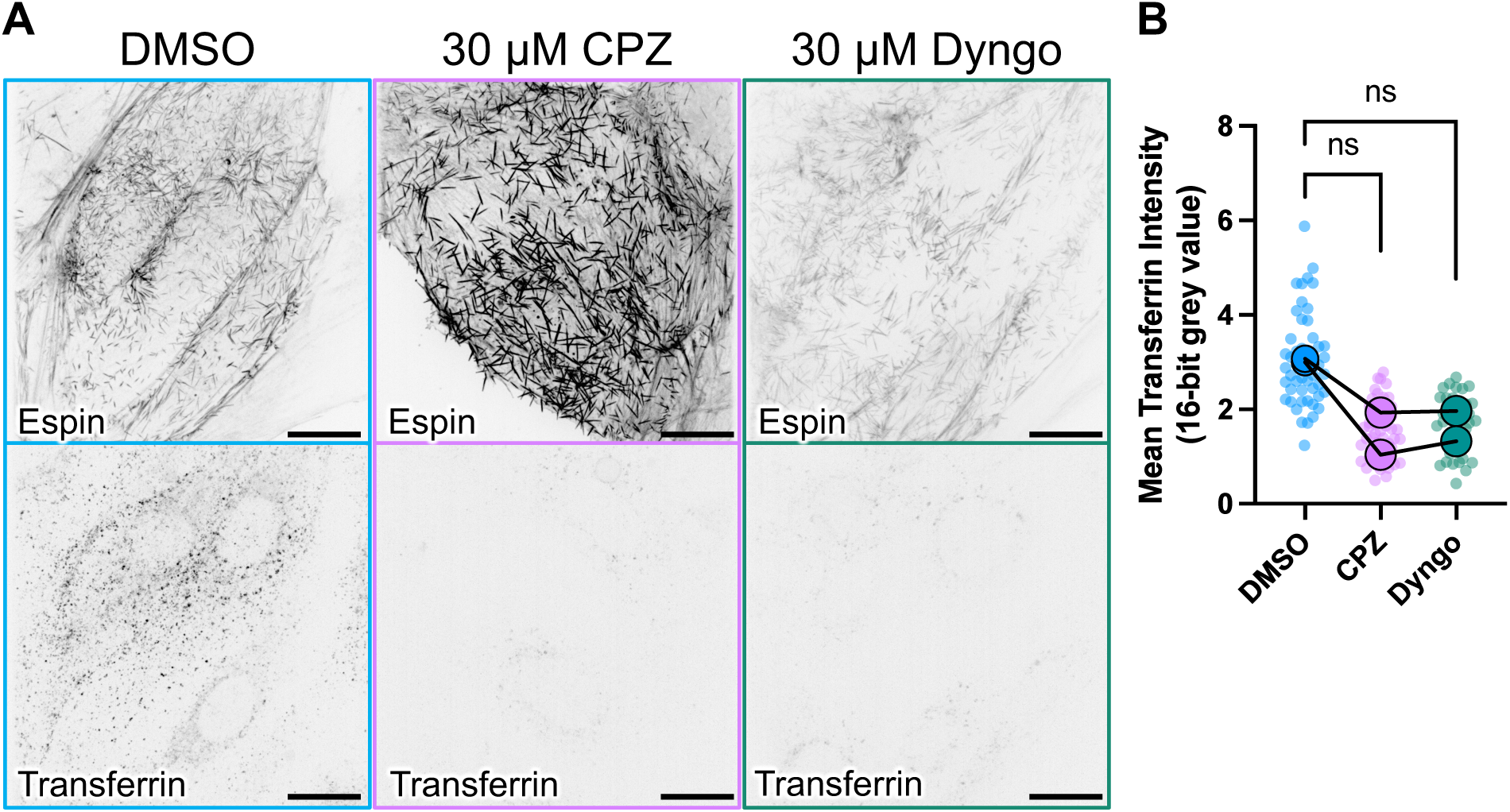
CPZ and dyngo equally inhibit transferrin uptake. (A) Maximum intensity projection (MIP) of CL4 cells transfected with mCherry-espin incubated with human purified transferrin after 45-minute treatment of DMSO, 30 µM CPZ, or 30 µM Dyngo. Scale bar = 10 µM (B) Superplot quantification of whole cell mean transferrin intensity measurements. Small transparent points represent individual cells, larger opaque points show experimental replicates. n = 100 cells sampled from two experimental replicates.

**Figure S3.**
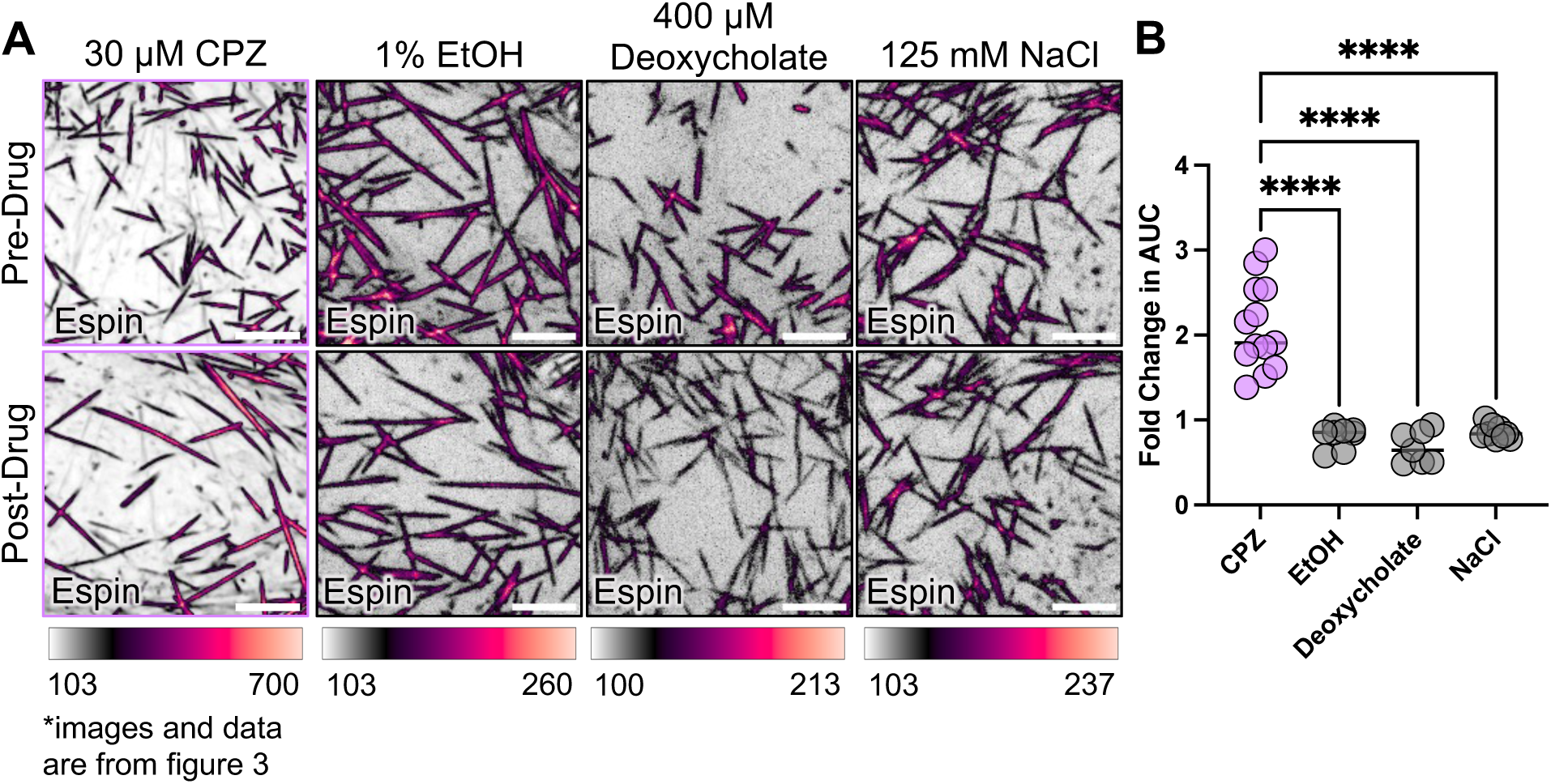
Global reduction in plasma membrane tension does not lead to the formation of exaggerated microvilli. (A) Confocal maximum intensity projection (MIP) of representative CL4 cells before and after treatment with CPZ, 1% EtOH, 400 µM Deoxycholate, and 125 mM NaCl. Cells were treated with CPZ and Deoxycholate for 45 minutes, and with 1% EtOH and 125 mM NaCl for 5-10 minutes. Drug was added immediately after 0 min image was captured. Single channels show intensity-coded color profile. Intensity scales from low (white) to high (yellow). Scale bar = 5 µm. (B) Fold change in area under the curve of mCherry-espin signal of microvilli post- vs. pre- drug treatments. 8-10 cells from 2 experimental replicates, 100-150 microvilli per condition. Images and data for the 30 µM CPZ treatment were taken from figure 3.

**Supplemental Figure 4.**
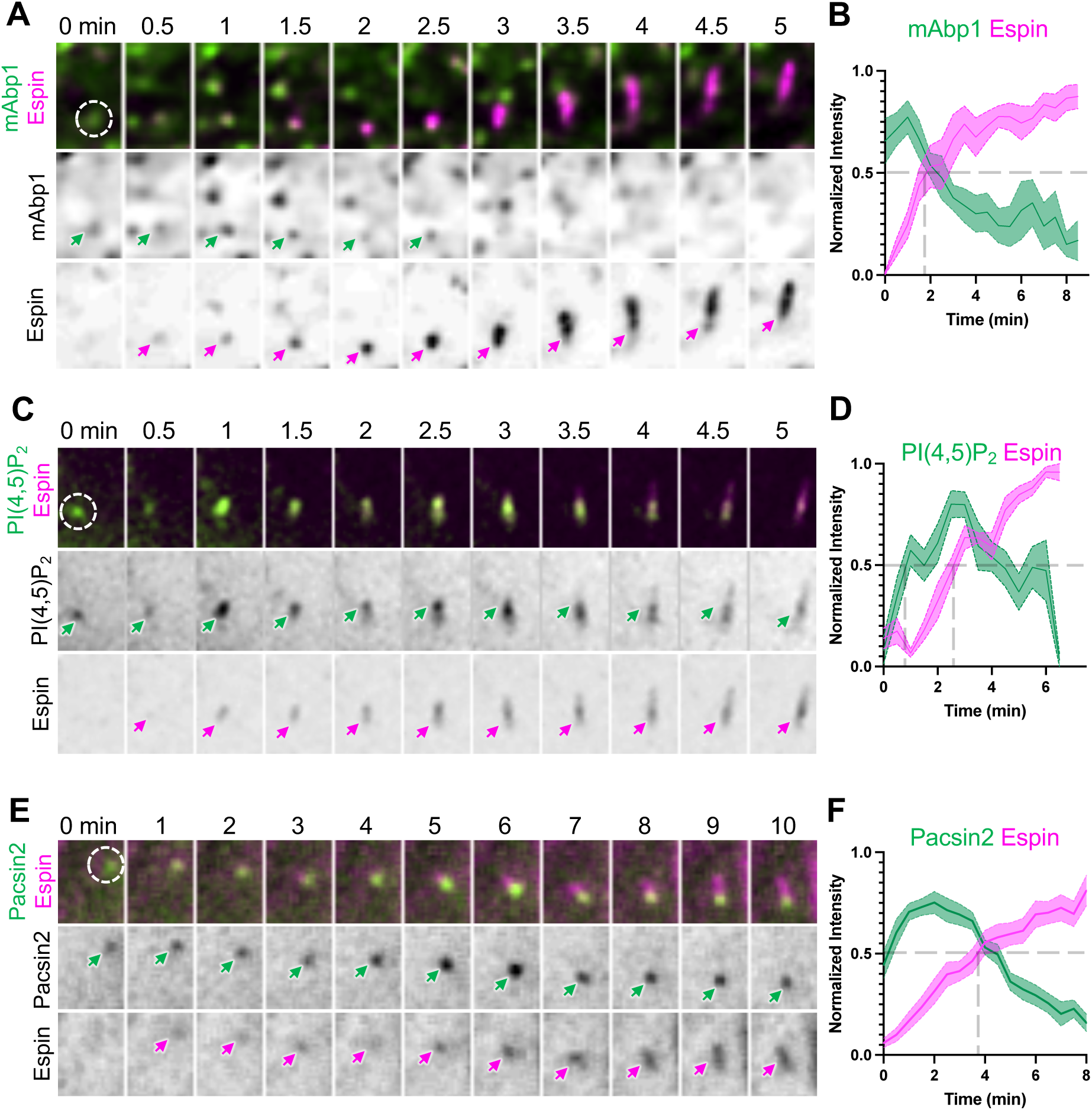
Microvilli grow from apical clathrin coated pits. (A) Montage of a single microvillus growth event in CL4 cells expressing EGFP-mAbp1 and mCherry-Espin. Every column represents 30 seconds. Arrows mark the analyzed core bundle. Column width = 2.09 µm. (B) Average fluorescence intensity measurements over time. n = 9 growth events from 4 cells across 2 independent experiments. (C) Montage of a single microvillus growth event in CL4 cells expressing EGFP-pPLC-ΔPH-GFP (PIP(4,5)P_2_ marker) and mCherry-Espin. Every column represents 30 seconds. Arrows mark the analyzed core bundle. Column width = 2.09 µm. (D) Average fluorescence intensity measurements over time. n = 20 growth events from 10 cells across 3 independent experiments. (E) Montage of a single microvillus growth event in CL4 cells expressing EGFP-pacsin2 and mCherry-espin. Every column represents one minute. Arrows mark the analyzed core bundle. Column width = 2.09 µm. (F) Average fluorescence intensity measurements over time. n = 20 growth events from 10 cells across 3 independent experiments.

**Figure S5.**
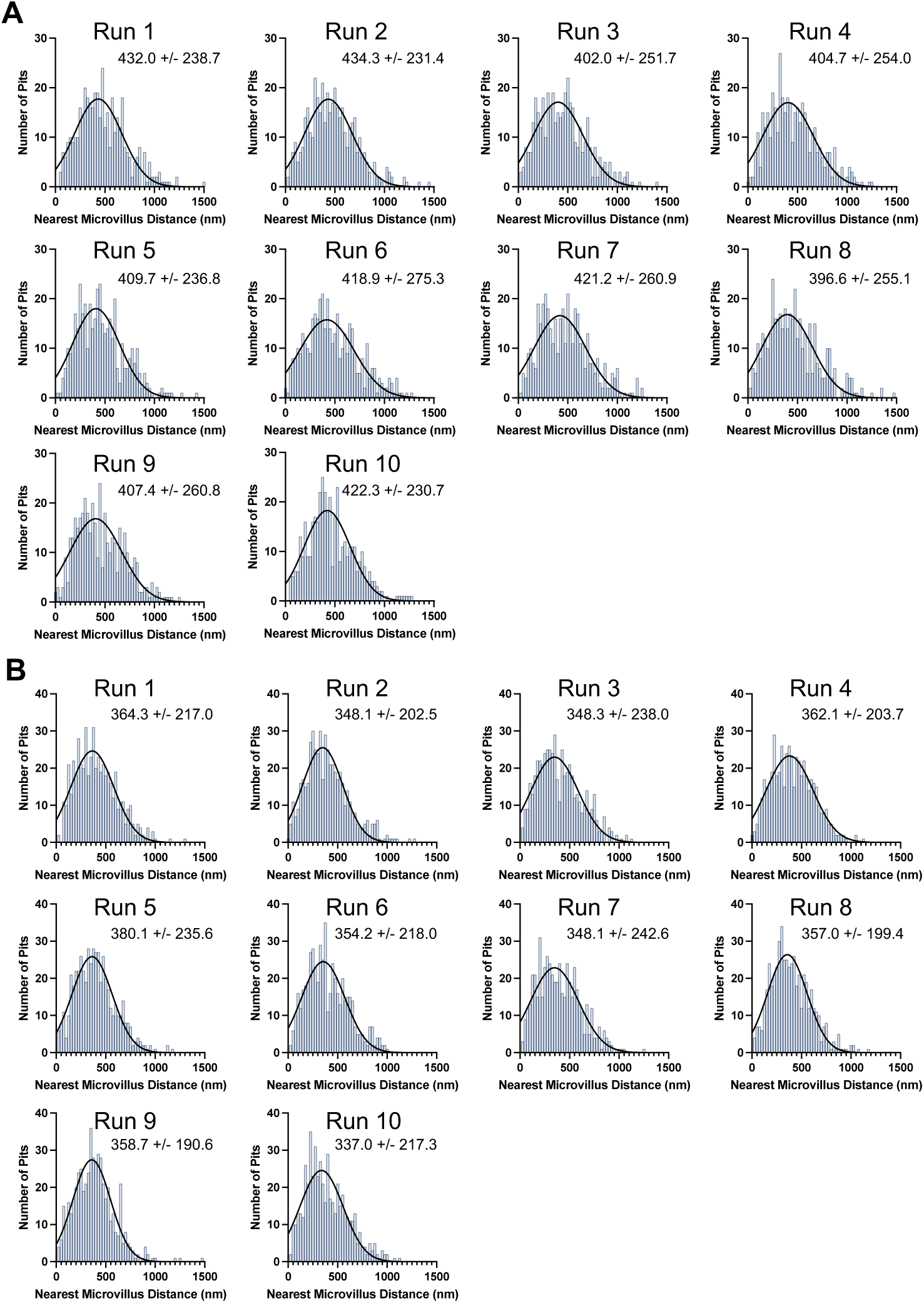
Simulation of randomly distributed endocytic pits and microvilli leads to nearest neighbor distances with gaussian distributions. (A) 10 runs of simulated nearest neighbor distance measurements from endocytic pits to nearest microvillus on the surface of CL4 cells. Simulated densities were matched with experimental measurements of 516 pits, 2570 microvilli over 1601.25 µm^2^. (B) 10 runs of simulated nearest neighbor distance measurements from endocytic pits to nearest microvillus on the surface of CACO_2_-_BBE_ cells. Simulated densities were matched with experimental measurements of 415 pits, 2039 microvilli over 1828.7 µm^2^.

